# Genome-wide analysis reveals distinct genetic mechanisms of diet-dependent lifespan and healthspan in *D. melanogaster*

**DOI:** 10.1101/153791

**Authors:** Kenneth A. Wilson, Christopher S. Nelson, Jennifer N. Beck, Rachel B. Brem, Pankaj Kapahi

## Abstract

Dietary restriction (DR) robustly extends lifespan and delays age-related diseases across species. An underlying assumption in aging research has been that DR mimetics extend both lifespan and healthspan jointly, though this has not been rigorously tested in different genetic backgrounds. Furthermore, nutrient response genes important for lifespan or healthspan extension remain underexplored, especially in natural populations. To address these gaps, we utilized over 150 DGRP strains to measure nutrient-dependent changes in lifespan and age-related climbing ability to measure healthspan. DR extended lifespan and delayed decline in climbing ability on average, but there was no evidence of correlation between these traits across individual strains. Through GWAS, we then identified and validated *jughead* and *Ferredoxin* as determinants of diet-dependent lifespan, and *Daedalus* for diet-dependent physical activity. Modulating these genes produced independent effects on lifespan and climbing ability, further suggesting that these age-related traits are likely to be regulated through distinct genetic mechanisms.

## Introduction

Dietary restriction (DR), the reduction in total nutrients (1, 2) or specific macromolecules (3–6) without malnutrition, is a robust method shown to extend lifespan and slow age-related dysfunction in multiple species. Genes mediating responses to dietary change are of prime interest in the development of anti-aging therapeutics (7), but surprisingly few are known. Conservation of signaling pathways and the rapidity with which lifespan studies can be carried out make research in model organisms critical to understand the molecular basis of aging and age-related diseases in humans (8–13). Some well-characterized nutrient-response pathways including the target of rapamycin (TOR) (14) and insulin-like signaling (ILS) (15) as well as sirtuins (14) have been proposed to mediate the effects of DR in species as diverse as yeast, worms, flies, and mice (16–22). However, these and other DR response pathways have largely been found through candidate-based screens. It is not clear whether these pathways are critical for conferring the diet-dependent changes in natural populations. Thus, there is an urgent need in the field to undertake whole-genome-scale studies in natural populations of multicellular organisms. In *D. melanogaster*, by varying the two components of the media, yeast extract (the primary source of protein and lipids) and sucrose, we and others have shown that nutrient composition and not just total caloric intake modulates metabolism, healthspan, and lifespan (23–25). Likewise in mice and humans, a low-protein, high carbohydrate diet has the maximal benefit of extending healthspan (3, 26). However, some animal studies have challenged the universality of the benefits of DR (27–29). A compelling explanation for these discrepancies is that natural genetic variation may influence the response to dietary change (28). There is evidence in humans that the selective pressures on the response to nutrient availability may vary across populations, resulting in natural genetic differences that may influence diabetes and obesity (30). Consistent with this notion, recombinant inbred strains of mice differ widely in their response to caloric restriction (31, 32), but the mechanisms behind this phenomenon are not understood and could be affected by natural genetic variation. Though the latter studies of DR effects across wild individuals were initially met with great enthusiasm in the field, the genes responsible for this variation have yet to be identified.

Measuring lifespan in response to DR has been the gold standard to identify the mechanisms that mediate the protective effects of DR in invertebrates (33, 34). An underlying assumption in the field is that if an intervention slows the rate of aging, then it would extend lifespan and also other healthspan traits (35). However, measuring lifespan has limitations as a measure of aging, and thus it is imperative also to assess healthspan to find the most promising interventions for humans (36, 37). Recent studies in humans have investigated the period at the tail end of life and how it correlates with mortality, and results have demonstrated that although disability generally increases as individuals approach death (38), the rate and severity of decline varies by case and individual (39). Studies in worms (40–43) and mice (36, 44) in select genetic backgrounds demonstrate that lifespan extension is not necessarily accompanied by an increase in healthspan. Thus these studies pose the question whether lifespan and healthspan are indeed determined by the same genetic mechanisms. Walking speed in humans, a strong indicator of health, is known to decline with age and is a predictor of mortality (45–48). Flies have an innate tendency to climb upwards in their enclosure. This climbing ability also declines with age and is widely used as a measure of healthspan (49–54). Previous studies on measuring healthspan have utilized interventions that are known to extend lifespan to examine the relationship lifespan and healthspan (35, 55), but mechanisms for healthspan extension in models without lifespan-dependent effects have not been examined. Thus, there is a need to examine the relationship between lifespan and healthspan in an unbiased manner in diverse genetic backgrounds.

Genome-wide association studies (GWAS) have become the standard to determine novel genetic regulators of longevity or health in humans (56–58) and model organisms (59–71), yet it has not been used to determine how diet impacts these complex traits. To fill this gap, we have used a genome-scale dissection of nutrient-responsive effects on lifespan and age-related climbing ability in the fly, using as a tool the genetic variation present in wild fly populations. As our library of genetic diversity, we use the *Drosophila* Genetic Reference Panel (DGRP) (72), a population of 205 genetically distinct wild fly lines, which has been established by the Mackay lab. These lines have been successfully used for GWAS of dozens of traits, including courtship songs (73), endoplasmic reticulum stress response (74), olfaction (75), and susceptibility to viral infection (76), as well as fecundity and lifespan in flies, reared on a single diet (59). The fly model offers the opportunity for an unbiased examination of the relationship between healthspan and lifespan. Identifying genes related to these traits will provide a better understanding of the genetic architecture that optimizes lifespan and healthspan in response to dietary interventions.

Using the DGRP collection, we examined the diet-dependent changes in lifespan and climbing ability across ages. We observed significant variation in the diet-dependent changes in both climbing ability and lifespan across the assayed DGRP lines. We have used GWAS to discover novel genetic regulators of longevity and health. Our results failed to demonstrate any significant correlation between these two phenotypes across the strains. We also identify several genetic variants that determine either physical functionality or longevity in a diet-dependent manner. We validated the effects of *CG34351*, which we name “*jughead*” (*jgh*), and *Fdxh* for diet-dependent changes in lifespan. We also validated that *CG33690*, a previously uncharacterized gene, has a role in diet-dependent changes in physical activity, and thus propose the new name “*Daedalus*” (*dls*). In addition to ascribing novel functions to these genes, we observe that these genes independently determine lifespan or age-related climbing ability but not both together.

## Results

### Genetic variation in diet-dependent changes in lifespan

To determine the effects of genetic variation on DR-mediated changes in lifespan, we reared ∼200 non-virgin females in eight vials from 161 DGRP lines on two dietary conditions that featured a ten-fold variation in dietary yeast. Our DR condition contained 0.5% yeast extract and *ad libitum* (AL) diet, 5% yeast extract as described previously (4, 77). We observed a broad range in diet-dependent changes in lifespan across the strains, ranging from a 65% reduction in median lifespan from AL to DR to a 12.5-fold increase (Fig. 1A). 83% of all lines survived longer on DR than on the AL diet (Fig. 1B). Out of the longer-lived DGRP strains on an AL diet (>35 days median lifespan), less than 46% received additional longevity benefits by DR, whereas 82% or shorter-lived strains (<35 days median lifespan) showed increased median lifespan when undergoing DR. We repeated lifespan measurements of 52 strains and saw largely reproducible median lifespans (Fig. 1C, R^2^ = 0.57 for AL, R^2^ = 0.64 for DR). In agreement with prior reports (28, 31, 78), we observed that DR extends lifespan in most, but not all, strains. We also observed that overall DR was less effective in extending the lifespan of strains that were already relatively long-lived.

**Figure 1.**
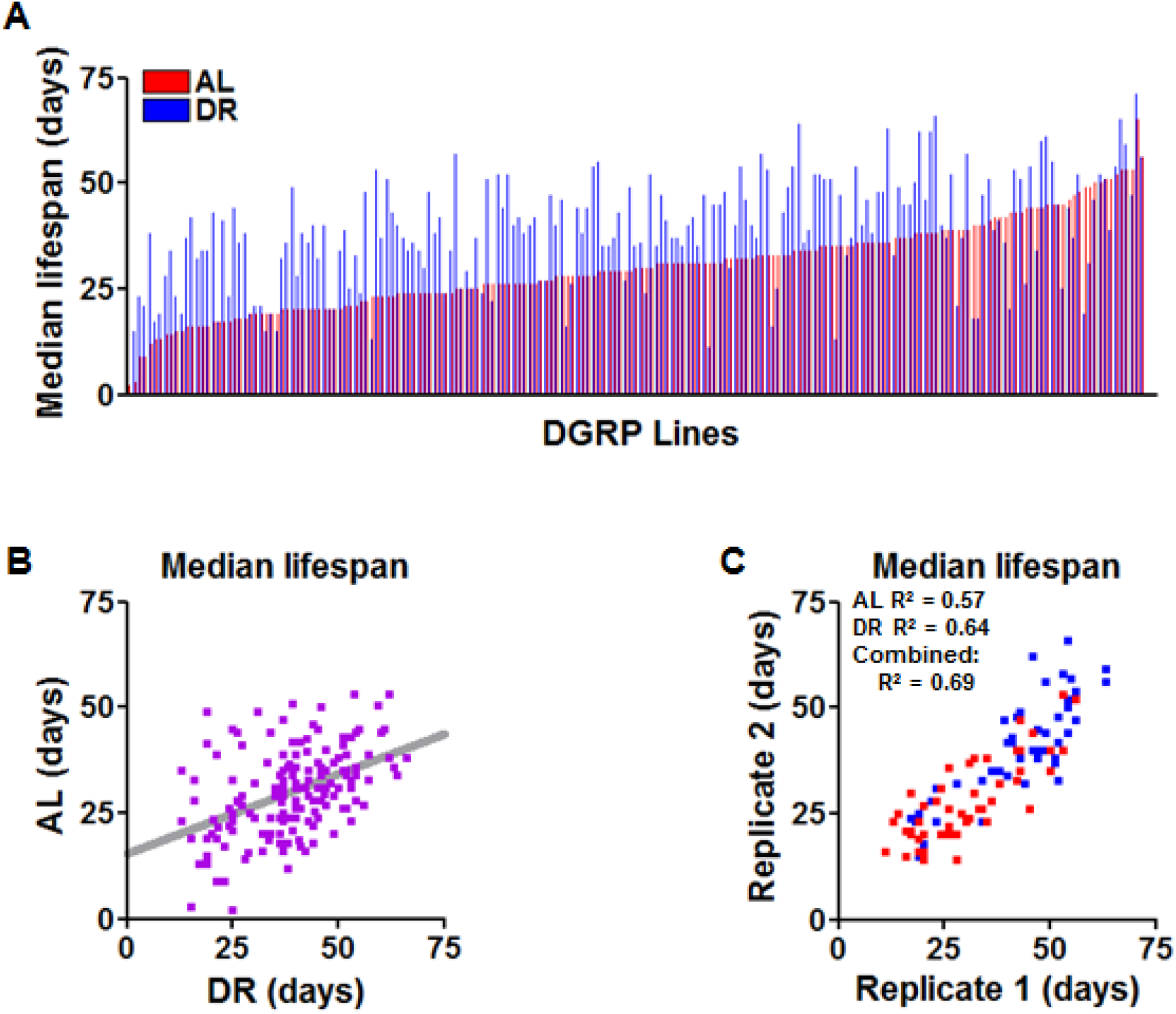
Genotype influences variation in lifespan and response to DR across the DGRP lines. (A) Median lifespan of 161 DGRP lines in ascending order on AL diet (red). Adjacent lines in blue represent the same strain raised on DR diet. (B) Comparison of median lifespan on AL of each strain with its DR counterpart. Same data as in A, displayed as a scatterplot. Grey bar represents best-fit trendline. (C) Comparison of median lifespan values across biological replicates of 52 DGRP lines on AL (red) and DR (blue). N = 200 flies per strain per diet.

### Decline in negative geotaxis with age varies by genotype and diet

In addition to measuring lifespan, we were interested in determining the genetic basis of functional health across the DGRP, as identifying means to alter health can substantially improve the quality of life. Because walking speed is frequently used as a marker of health and a predictor of mortality in humans (79, 80), we utilized flies’ natural tendency to climb their enclosure to track negative geotaxis performance in 156 DGRP lines throughout the duration of their life in parallel to measuring lifespan, as described above. We measured the percentage of each line able to climb an empty vial wall once per week throughout adulthood (81–83) (detailed in Methods). As an index of health decline, we analyzed the day at which a given DGRP strain fell below 50% of its initial climbing ability. Some strains fell below this threshold within the first week after being placed on DR or AL while others maintained greater than 50% climbing capacity for longer than 60 days (Fig. 2A). Consistent with previous reports, we found that DR generally improved physical activity. We observed that DR delayed the age-related decline in climbing ability in 69% of all tested lines, with another 25% of lines showing no difference between the two diets (Fig. 2B).

**Figure 2.**
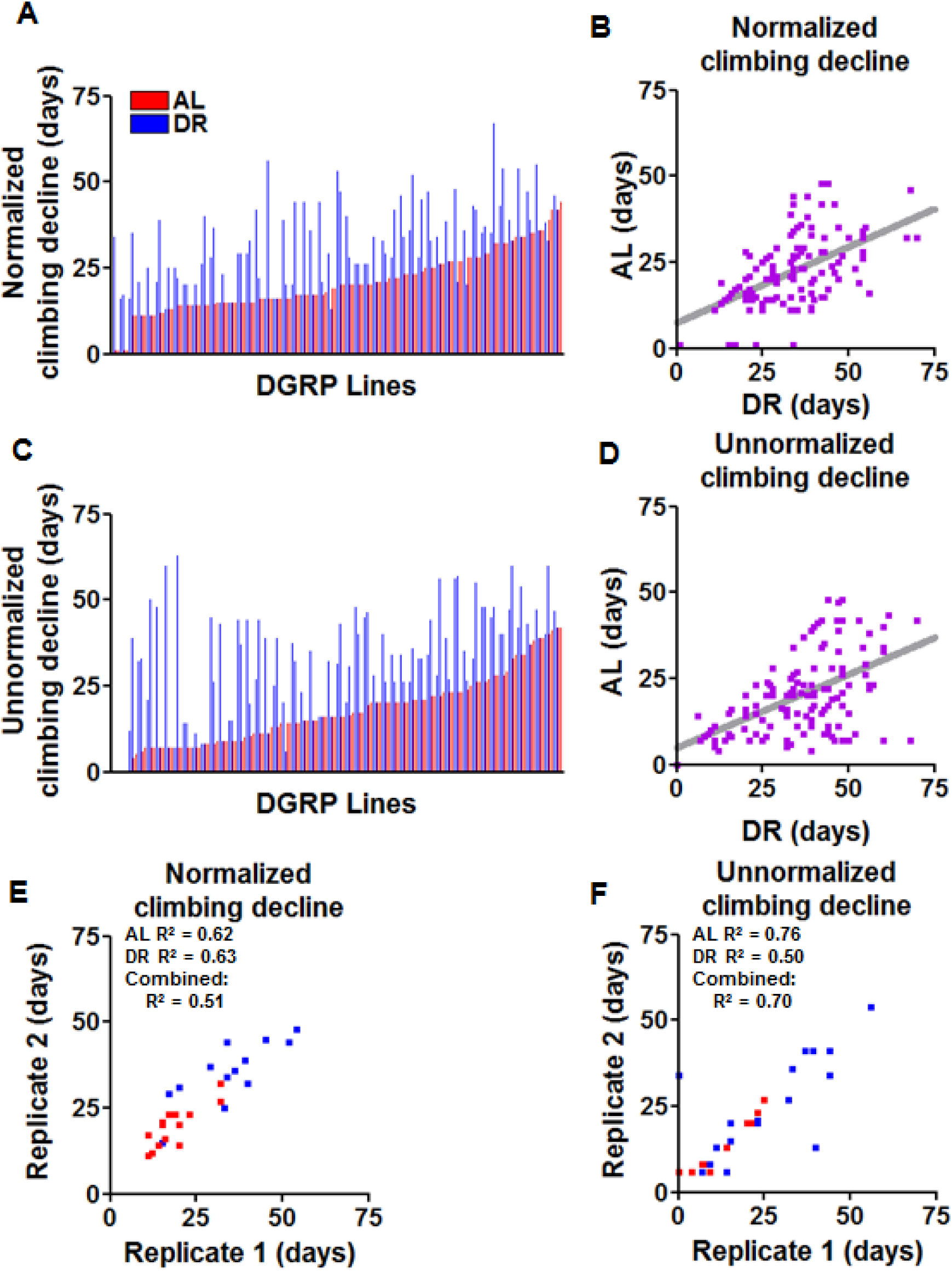
Decline in climbing ability varies by genotype. (A) The age (in days) at which a line declines to half of its initial percent of climbing flies. Data are arranged in ascending order of the strains’ AL phenotypes (red). Adjacent lines in blue represent the same strain on DR diet. Comparison of days below 50% maximal climbing capacity of each strain on AL versus DR diet. Grey bar represents best-fit trendline. (C) The age (in days) at which fewer than 20% of the surviving population can climb in the allotted time. Data are arranged in ascending order by the phenotype on AL diet (red) with adjacent lines representing the same strain on DR (blue). (D) Comparison of each strain’s climbing data between AL and DR diets. Grey line represents best-fit trendline. (E) Comparison of biological replicates of 25 tested DGRP lines on AL (red) or DR (blue) for 50% decline in initial climbing ability. (F) Comparison of biological replicates for 25 tested DGRP lines for the day at which less than 20% of surviving flies are able to climb on AL (red) and DR (blue). N = 200 flies per strain per diet.

Since normalizing to initial climbing ability removed the variation in the absolute climbing ability, we also analyzed the changes in the absolute percentage of flies that were capable of climbing across strains. For this trait, we recorded the day at which the percentage of surviving animals able to climb fell under 20% (Fig. 2C, Suppl. Fig. 1). DR extended the length of time above 20% climbing ability over AL in 87% of lines, with 12% of lines declining at the same day of life regardless of diet (Fig. 2D). For both climbing measures we re-tested 17 lines and found our recorded values to be reproducible (Fig. 2E-F, R^2^ = 0.62 for AL 50% decline and 0.63 for DR, R^2^= 0.76 for AL day below 20% climbing and 0.50 for DR). Together, these results indicate that DR generally improves climbing ability, but the degree to which it is beneficial varies by genotype.

### Lack of correlation between recorded lifespan and climbing values

To better understand the relationship between healthspan and lifespan, we compared the relationship between median lifespan and age-related climbing ability. Separating the strains on AL into the longer-lived half of strains (>35 days median lifespan) and shorter-lived (<35 days), we found that the average day of Across the shorter-lived half of the strains on the AL diet (<35 days median lifespan) the average median lifespan was 23.9 days and the average day these strains reach half of their initial climbing ability was 19.4, meaning on average these strains maintained better than half their initial climbing ability for 81.2% of their average median lifespan. Across longer-lived strains on AL conditions (>35 days median lifespan), the average median lifespan was 38 days and the average normalized day of climbing decline was 22.9, 60% of the average median lifespan (Fig 3A). On the DR diet, the shorter-lived strains (<39 median lifespan) had an average median lifespan of 30.7 days and average day of climbing decline 28.2, maintaining climbing ability for an elevated 91.9% of their lives. In long-lived strains (>39 days median lifespan), the average median lifespan was 46.8 days and the average day of climbing decline was 33.9, or only 72.4% of the average median lifespan (Fig. 3B). Looking deeper into these phenotypes across the individual strains, we found no evidence of a correlation between median lifespan and 50% climbing decline on the AL across individual strains (Fig. 3C, R^2^ = 0.05), nor the DR diet (Fig. 3D, R^2^ = 0.07). Similarly, we found no correlation between median lifespan and our absolute climbing decline value (Suppl. Fig. 2, AL R^2^ = -0.06, DR R^2^ = -0.01). We also looked into responsiveness of each DGRP strain to DR, and found that across all of the strains, only 50% of all strains showed >3 days improvement in both lifespan and days of life above 50% initial climbing capacity in response to DR. Alternatively, 14% of strains showed opposing phenotypes, either with reduced climbing ability and increased lifespan on DR, or vice-versa. The remaining 36% showed no change in either or both phenotypes (Fig. 3E). We found no evidence of a correlation between change in climbing ability and change in median lifespan in response to DR (Fig. 3F, R^2^ = -0.04). Together, these results imply that though DR overall extends lifespan and healthspan, when examined across the individual DGRP strains these two traits fail to correlate in our data.

**Figure 3.**
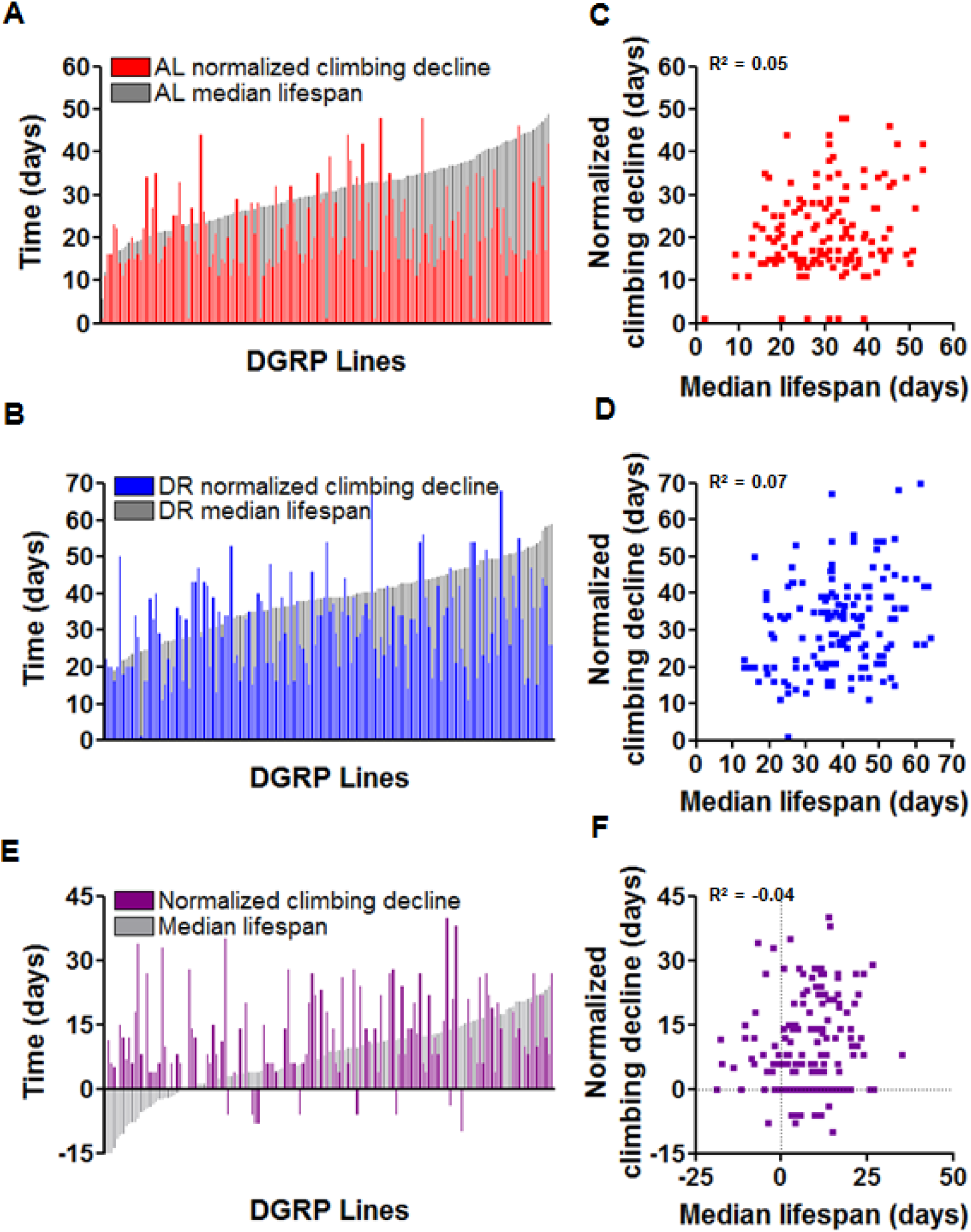
Genotype and diet differentially influence lifespan and healthspan separately. (A and B) Comparison of each tested strain’s day below 50% of maximal climbing proportion with median lifespan, on (A) AL or (B) DR. Each bar represents a DGRP strain, ordered by median lifespan on each diet. Colored bars represent climbing half-life and white bars represent median lifespan. (C and D) Scatter plots depicting climbing ability compared to median lifespan on the AL diet or (D) DR. Each dot represents a single DGRP strain. (E) Comparison of DR responsiveness with regards to median lifespan (white bars) and time above 50% initial climbing ability (purple bars). (F) Scatter plot depicting response to DR of each tested DGRP line with regards to median lifespan and amount of time above 50% initial climbing ability. Each dot represents a single DGRP strain. N = 200 flies per strain per diet.

### Genome-wide association analysis

Next, we determined the genetic basis for the phenotypic differences in median lifespan and climbing ability across the DGRP. We performed genome-wide association studies (GWAS) using a linear regression model with terms for genotype, diet, and the interaction between genotype and diet as described in the Methods (called “Interaction” terms). We identified a list of candidate loci with a minor allele frequency ≥25% with statistical signals of ≤10% FDR based on permutation analysis (Table 1, Methods). Included among the candidates for lifespan regulation, indicated in Table 1, were variants in the genes *CR32111* (three variants), *CG43203* (one variant), *jgh* (one variant), and *CG8312* (one variant). These variants were significantly associated with regulating diet-dependent changes in the day at which 75% of a population was surviving, determined through our the interaction term in our GWAS model. As regulators of median lifespan, we identified variants in *CG5888* (one variant), *CR32111* (three variants, same three as associated with 75% survival), *CG31221* (one variant), and *CR45580* (one variant) in interaction with diet. GWAS also identified a variant in *CG5888* as regulating longevity in a diet-independent manner based on genotype alone (Table 1).

**Table 1.**
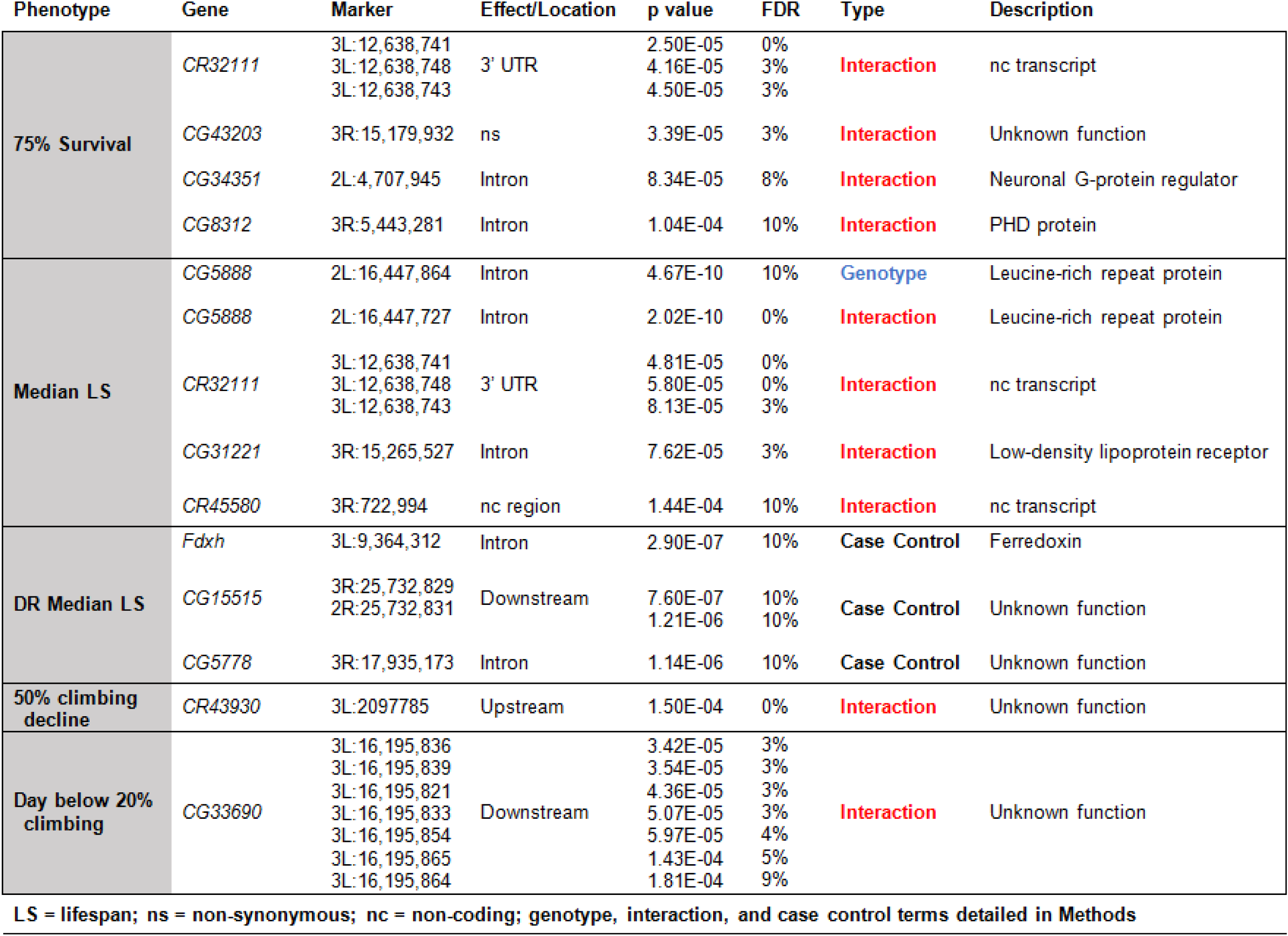
Lifespan and climbing gene candidates identified by genome-wide association.

We next searched for loci that associated with exceptional longevity (see Methods) as an additional screen for genetic variants that contributed to the extreme length of life rather than an increased median lifespan. For this, we determined the upper 15^th^ percentile of longevity across all tested strains (>41 days median lifespan on AL and >51 days on DR) as our “long-lived” lines. This restructured “Case/Control” GWAS detected an association between DR longevity and a polymorphism in *Ferredoxin 1* (*Fdxh*) as well as two variants downstream of *CG15515* and one intronic variant in *CG5778* (Table 1).

To further examine the relationship between lifespan and healthspan, we determined the genetic loci associated with changes in climbing ability. GWAS for the day of 50% climbing decline from initial climbing ability identified one significant polymorphism upstream of the non-protein coding gene *CR43930* (Table 1). For the day at which fewer than 20% of surviving flies could climb, GWAS identified seven polymorphisms downstream of *dls* to be significantly associated with climbing in a diet-dependent manner (Table 1).

### Diet and tissue-specific changes in *dls*, jgh, and *Fdxh*

We conducted a preliminary RNAi screen with all of our candidate genes to determine how altered candidate gene expression could impact longevity or climbing ability. With the use of the whole-body expression driver *Act5C*-GS, we induced RNAi in five of the candidate genes indicated through GWAS (Suppl. Fig. 3). We found that whole-body RNAi of *CR32111* resulted in a 10% reduction in median lifespan on DR but no change on AL (Suppl. Fig. 3A). RNAi of *CG8312* resulted in a slight, 3% reduction of median lifespan on DR and a 7% extension in life on AL (Suppl. Fig. 3B). RNAi of *CG5888* resulted in no change to median lifespan on DR and no change on AL (Suppl. Fig.3C). RNAi of *CG31221* and observed a 26% reduction in median lifespan on DR and no change on AL (Suppl. Fig. 3D). RNAi of *CG15515* resulted in a 5% reduction of median lifespan on DR and no change on AL (Suppl. Fig. 3E). Based on FlyAtlas data, we induced RNAi of *CG5778* in the fat body with the *S106*-GS-Gal4 and observed 5% reduction of median lifespan on DR and a 13% reduction in median lifespan on AL (Suppl. Fig. 3F). We chose to focus further on our three other candidate genes, *dls, Fdxh*, and *jgh*, to determine their role in modulating diet-dependent changes lifespan and healthspan.

Of the seven loci downstream of *dls* associated with climbing regulation (Table 1), the most significant was found on chromosome 3L at position 16,195,836. At this locus, strains with a G allele showed significant delay in climbing decline over those with a T allele upon DR, but no difference was noted under AL conditions (Fig. 4A-B). We examined tissue specificity and mRNA expression changes in response to diet to aid in understanding the mechanism by which diet-dependent changes in phenotypes are mediate. We have previously generated data showing tissue-specific changes in mRNA translation state upon DR (33). With the use of FLAG-tagged ribosomal protein RPL13A, we pulled-down polysomes and analyzed the tissue-specific changes in mRNAs upon DR as previously described (33, 84). We observed that *dls* was moderately elevated in the germline, heart, and muscle (Suppl. Fig 4A). To test for transcriptional expression changes, we performed qRT-PCR for *dls* in body segments of *w*^*1118*^ control flies raised on either AL or DR diet for 7 days and saw elevated expression in the abdomen relative to other body segments as well as a nine-fold increase in expression in the head on DR versus AL diet (Fig 4C). As there were no RNAi constructs available for this line, we used a line containing a *Minos* element insertion in *dls* (Suppl. Fig. 9) (85) for our validation experiments. With this mutant line, we observed a ∼90% reduction in mRNA expression in DR conditions (Suppl. Fig. 5A-B). We saw a 19% increase in median lifespan on DR and 8% on AL in a *w*^*1118*^ background over *w*^*1118*^ controls (Fig. 4D) but a 7% reduction on DR and a 19% reduction in median lifespan on AL in a Canton-S background (Suppl. Fig. 6A). We found that climbing ability was consistently increased in mutants fed DR over their wildtype controls regardless of strain background, with no significant changes observed in AL conditions (Fig. 4E, Suppl. Fig 6B). In total, while the longevity effects of this mutation were mixed depending on strain background there was a significant improvement in climbing ability only observed on a DR diet (Fig. 4F, Suppl. Fig. 6C). We also tracked spontaneous activity for these flies for 24 hours (see Methods), and found that the *dls* mutant had increased spontaneous activity only on the DR diet regardless of strain background (Suppl. Fig. 8). Due to its role in regulating climbing and spontaneous activity on DR, we propose the common name *Daedalus* (*dls*) for *CG33690* after the mythological Greek inventor who created wings to escape incarceration by King Minos. Thus *dls*, which was identified as a candidate that influences diet-dependent changes in climbing ability, showed consistent effects on healthspan but failed to show consistent effects on lifespan. These data further argue that lifespan and age-related climbing ability are likely to be regulated by distinct mechanisms and that healthspan can be extended without a concomitant increase in lifespan.

**Figure 4.**
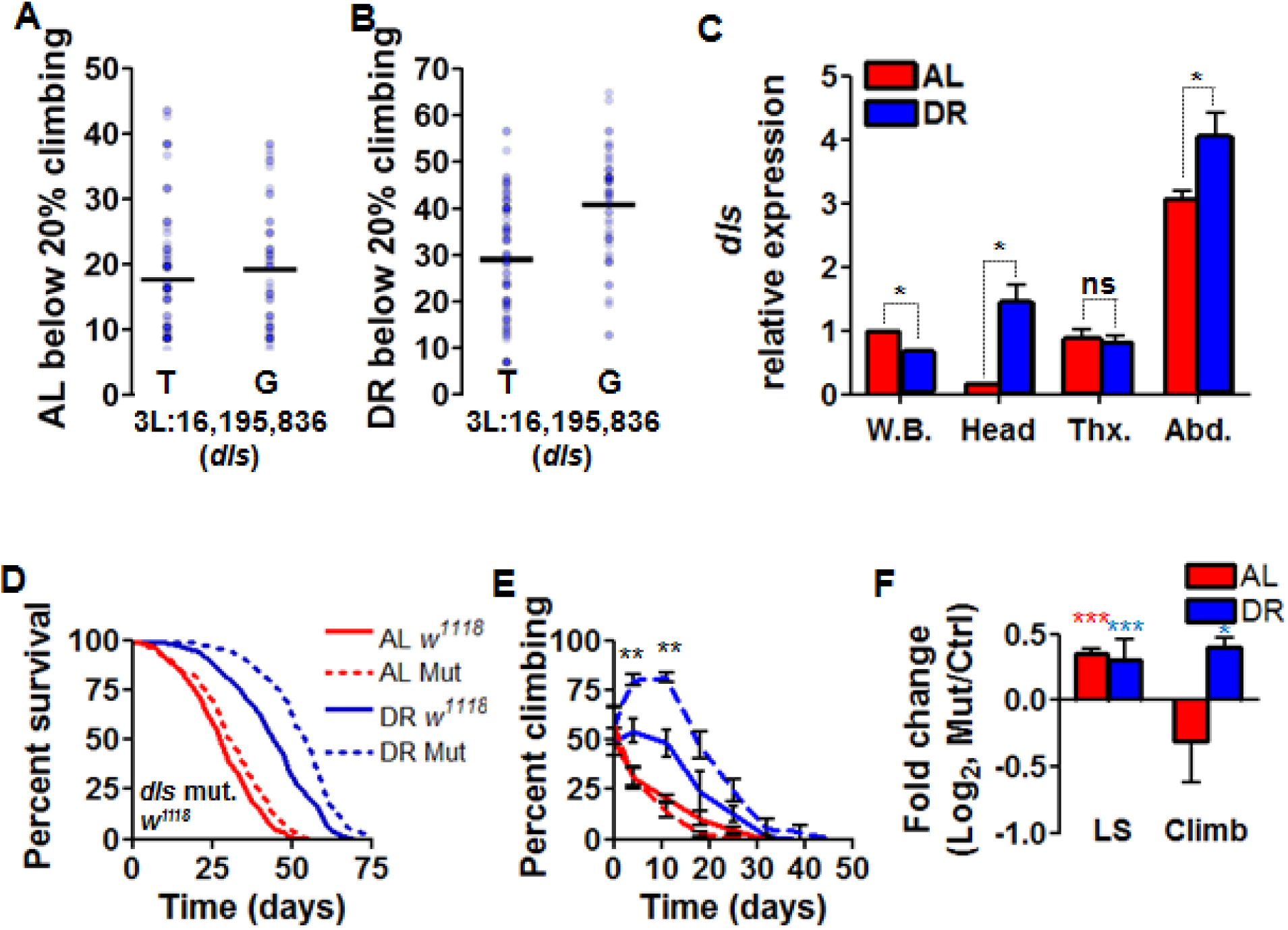
*Daedalus* regulates DR-specific climbing ability. (A and B) Plot of the day at which fewer than 20% of flies climb in tested DGRP lines, split by genotype at the most significant locus downstream of *dls* on (A) AL or (B) DR. p < 4E-5, FDR = 3%. (C) Expression of *dls* in the whole body (W.B.), head, thorax (Thx.), or abdomen (Abd.) of *w*^*1118*^ control strain after seven days of adulthood on AL (red) or DR (blue). The bars displayed are expression relative to the whole body AL divided by expression for *rp49* a housekeeping gene. N = 5 flies for whole body, 50 flies for heads, 50 thoraxes, and 50 abdomens repeated in biological triplicates. (D-F) The effect of *Minos* element insertion on (D) lifespan and (E) climbing ability over the course of life in a *w*^*1118*^ genetic background, and (F) the log_2_ difference median lifespan and unnormalized climbing decline between mutant and controls. AL shown in red, DR in blue. Significant differences between mutant and controls are indicated by *. * = p < 0.05, ** = p < 0.005, *** = p < 0.0005. nc = no change, ns = not significant. N = 200 flies per condition for each mutant experiment. Data in (D-I) show collective results from three biological replicates. Error bars represent SD between replicates.

Through our Interaction GWAS for longevity, we identified an intronic variant in *jgh* associated with the day at which less 75% of a strain’s population was surviving (Table 1). We found that DGRP strains fed the AL diet with a G allele at a particular locus (chr. 2L, position 2,707,945) showed a slightly delayed decline in 75% survival over strains with an A at that locus (Fig. 5A). Alternatively, strains with the G allele in DR conditions showed a significantly reduced 75% survival than counterparts with the A allele at the locus of interest (Fig. 5B). This gene is homologous to the human gene Regulator of G-Protein Signaling 7 Binding Protein (*RGS7BP*), and was previously noted in a GWAS screen for growth regulators of wing size (86) but has not previously been shown to play a role in longevity nor diet response. From our ribo-tag data we observed an eight-fold increase in *jgh* expression in the brain under DR conditions, as well as a 2.5-fold increase in expression on DR in the fat body and Malpighian tubule (Suppl. Fig. 4B). Through qRT-PCR we found that both on AL and DR diet *jgh* is expressed in the thorax, but DR induces a 55-fold increase in expression in the head (Fig. 5C). Thus, we used a pan-neuronal RU486-inducible *Elav*-GS-Gal4 driver to examine diet-dependent changes in longevity and health. We observed a 20% increase in median lifespan on an AL diet over control flies with one *jgh* RNAi strain (v30160, Suppl. Fig. 9, Fig. 5D) and a 29% increase in median lifespan on AL with a second strain (v30163, Suppl. Fig. 6D). We propose the name *jughead* (*jgh*) for this gene, after the fictional comic book character with a propensity for overeating without suffering its ill effects. In parallel, we examined whether inhibition of *jgh* will also extend healthspan. Surprisingly, RNAi of *jgh* failed to show any significant change in age-related climbing ability (Fig. 5E, Suppl. Fig 6E). Through qRT-PCR of the heads of flies from these crosses, we again saw elevated expression under DR conditions relative to flies raised on the AL diet, with RNAi-inducing approximately 50% reduction in the expression on AL and 70% reduction on DR. (Suppl. Fig. 5C-D). Overall we saw increased longevity on AL with *jgh* RNAi in neurons but did not see a change in the climbing ability with age (Fig. 5F, Suppl. Fig. 6F). Thus, inhibtion of *jgh* only extends lifespan without significant effects on climbing ability.

**Figure 5.**
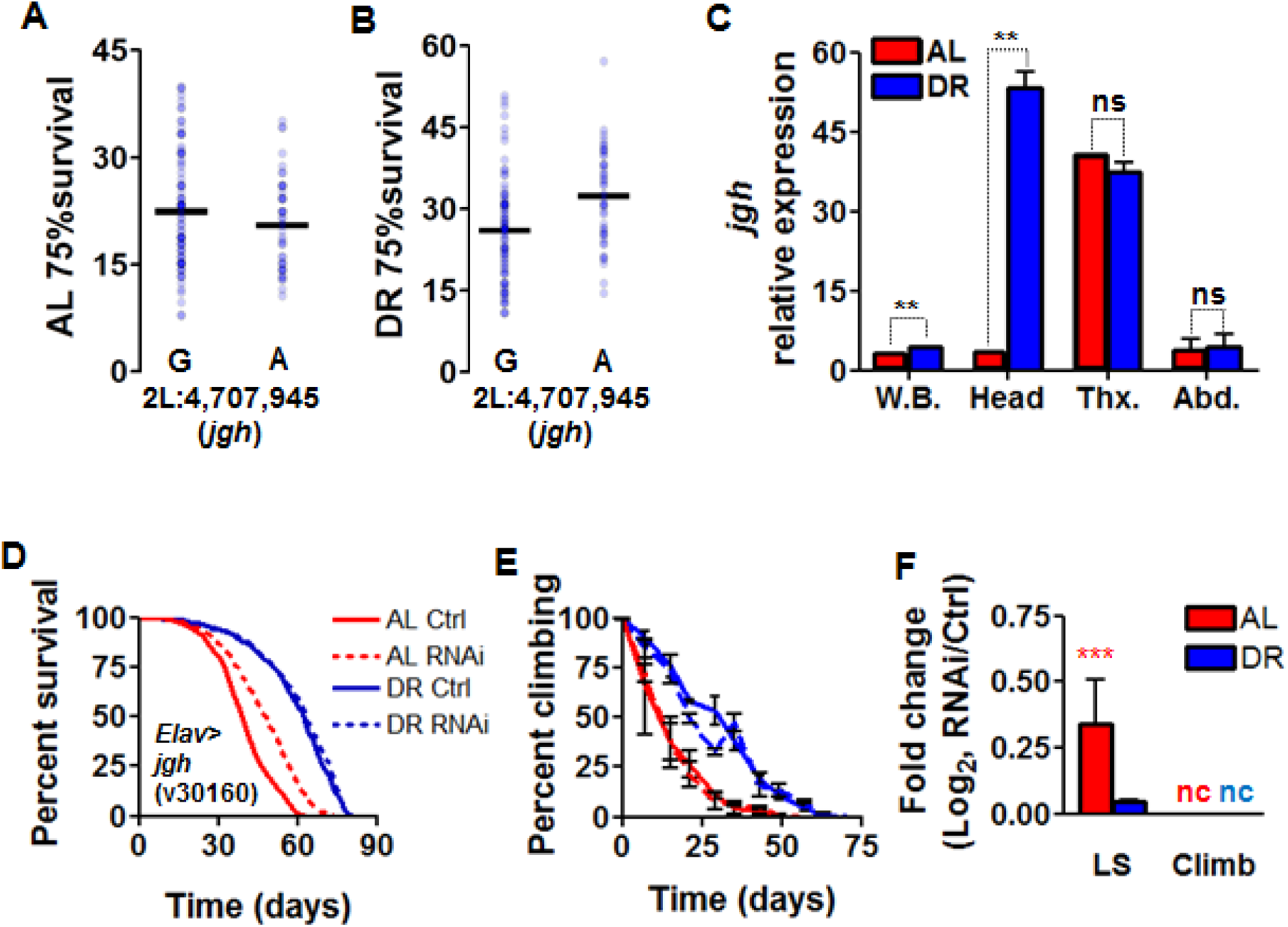
*jughead* regulates longevity diet-dependently. (A and B) Alignment of all 161 DGRP lines according to genotype at a particular locus in *jgh* and according to the day at which ≤ 75% of flies in a strain remain alive on (A) AL or (B) DR. Strains’ median lifespans are represented by blue dots, black bars represent mean values across all tested strains with a given genotype and diet. Significance for diet interaction p < 9E-5, FDR = 8%. (C) Expression of *jgh* in the whole body (W.B.), head, thorax (Thx.), or abdomen (Abd.) of *w*^*1118*^ control strain after seven days of adulthood on AL (red) or DR (blue). The bars displayed are expression relative to the whole body AL divided by expression for *rp49* a housekeeping gene. N = 5 flies for whole body, 50 flies for heads, 50 thoraxes, and 50 abdomens repeated in biological triplicates. (D-F) The effect of neuron-specific RNAi of *jgh* using the v30160 transgenic line in regulating (D) lifespan and (E) climbing ability over the course of life, with (F) log_2_ fold-changes between RNAi and control for both median lifespan and unnormalized climbing decline values. AL shown in red, DR in blue. Significant differences between RNAi and controls are indicated by *. * = p < 0.05, ** = p < 0.005, *** = p < 0.0005, determined by unpaired t test. nc = no change, ns = not significant. N = 200 flies per condition for each RNAi experiment. Data in (D-I) show collective results from three biological replicates. Error bars represent SD between replicates.

One locus identified through our Case-Control GWAS for strains with exceptional longevity (see Methods) was position 9,364,312 on chromosome 3L, which falls in an intronic region of the gene *Fdxh*. At this locus, DGRP strains with a C or A allele showed no difference in average median lifespan on AL (Fig. 6A), but under DR the A allele proved beneficial to median lifespan (Fig. 6B). *Fdxh* is involved in ecdysteroid production in flies (87), and human homologs have been implicated in mitochondrial maintenance (88, 89). Results from RPL13A tagging showed that under DR, *Fdxh* expression was modestly increased in every tissue except the heart and neurons (Suppl. Fig. 4C), and qRT-PCR results showed that DR induces moderately increased expression in each body segment (Fig. 6C). As such, we used *Act5C*-GS-Gal4 that leads to whole-body RNAi. Inhibition of *Fdxh* resulted in a 12% reduction in median lifespan on DR with the use of one *Fdxh* RNAi line (v104499, Suppl. Fig. 9, Fig. 6D) and a 20% decrease in median lifespan on DR as well as a 19% decrease on AL with the use of another RNAi line (v24497, Suppl. Fig. 9, Suppl. Fig. 6G). Using the transgenic line v1014499 we also saw no change in climbing ability at any point in life on either diet (Fig. 6E), but using v24497 resulted in a reduction in climbing ability in the second week of adulthood on DR (Suppl. Fig. 6H). In all, we found that whole-body RNAi of *Fdxh* causes a reduction in lifespan on DR and depending on the genetic background may also cause a reduction on AL, but the age-related climbing ability was unaffected (Fig. 6F, Suppl. 6I). qRT-PCR of whole body RNAi knockdown flies showed no change between DR and AL expression, with knockdown inducing a 50% reduction in expression (Suppl. Fig. 5E-F). Combined with our results from *dls* and *jgh* validation experiments, these results suggest a genetic uncoupling of lifespan and climbing phenotypes.

**Figure 6.**
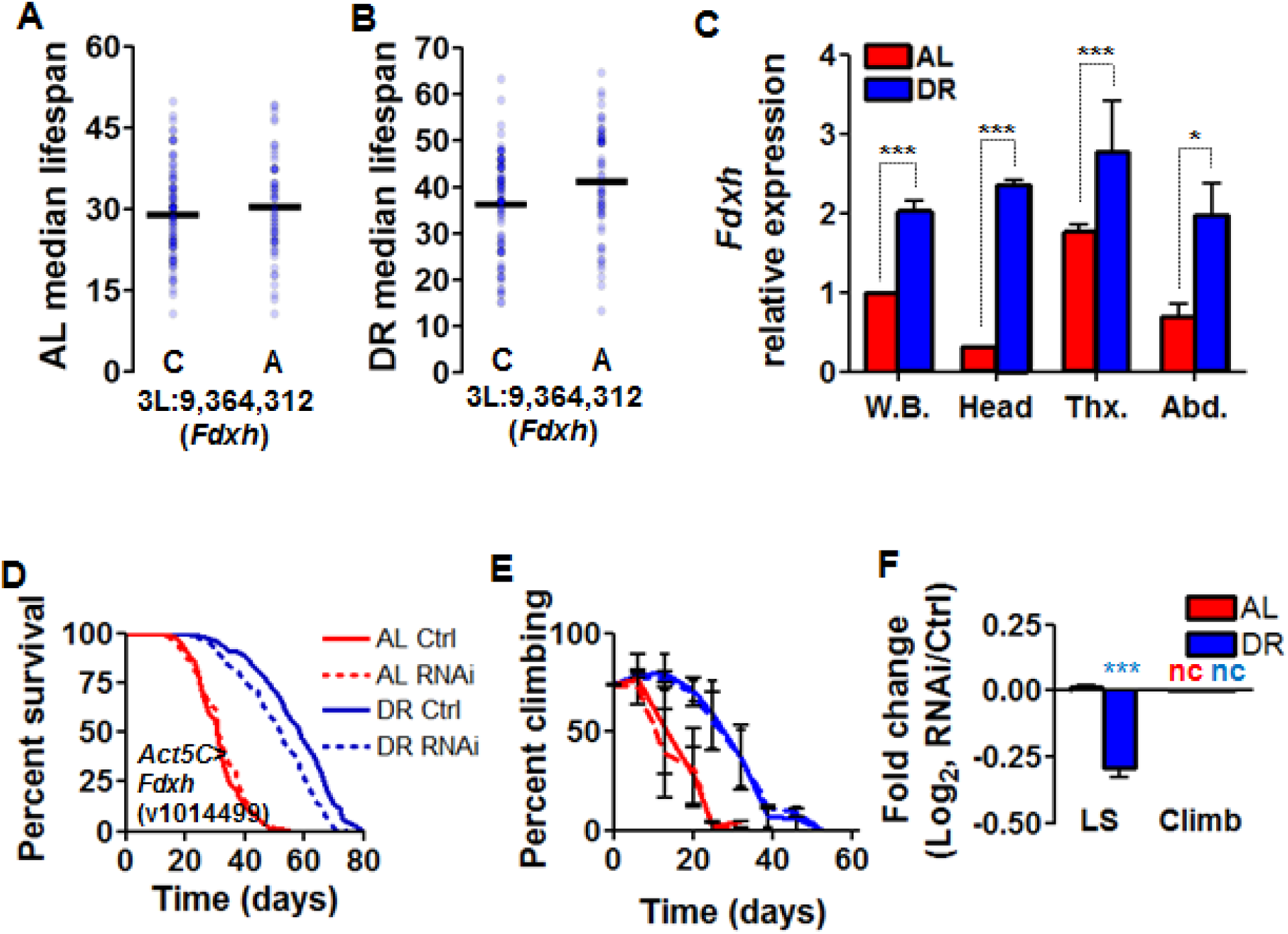
*Ferredoxin* regulates extreme longevity diet-dependently. (A and B) Alignment of all tested lines according to median survival at a particular locus in *Fdxh* on (A) AL or (B) DR. Difference in median lifespan on DR as determined by Fishers exact test, p < 3E-7, FDR = 10%. (C) Expression of *Fdxh* in the whole body (W.B.), head, thorax (Thx.), or abdomen (Abd.) of *w*^*1118*^ control strain after seven days of adulthood on AL (red) or DR (blue). The bars displayed are expression relative to the whole body AL divided by expression for *rp49* a housekeeping gene. N = 5 flies for whole body, 50 flies for heads, 50 thoraxes, and 50 abdomens repeated in biological triplicates. (D-F) The effect of whole-body RNAi of *Fdxh* using the v24497 transgenic line in regulating (D) lifespan and (E) climbing ability over the course of life, with (F) log_2_ fold-changes in median lifespan and unnormalized climbing decline between RNAi and controls represented. AL shown in red, DR in blue. Significant differences between RNAi and controls are indicated by *. * = p < 0.05, ** = p < 0.005, *** = p < 0.0005, determined by unpaired t test. nc = no change. N = 200 flies per condition for each RNAi experiment. Data in (D-I) show collective results from three biological replicates. Error bars represent SD between replicates.

## Discussion

Dietary restriction remains one of the most robust methods for lifespan extension and health improvement. Its benefits are widespread, including improved responses in models of cancer (90, 91), neurodegeneration (92), and other age-related disorders (93–95). Despite these reported benefits, enthusiasm for DR has been tempered by the observation that model organisms of different genotypes respond differently to DR (32, 77), with some genotypes even showing a reduced lifespan (31) or worsened health (96). Thus, identifying the mechanisms that promote longevity but not healthspan in response to diet may not provide the most suitable targets for humans.

Here, we have utilized the DGRP to perform the first longevity and healthspan GWAS featuring two different dietary compositions. We found that DR was beneficial in 83% of strains’ median lifespan and 87% of strains’ climbing ability, while the remainder of strains showed no effect or negative effects. These results are consistent with previous findings in flies and mice, where different nutrient manipulations failed to induce universal effects across strains (97, 98). Our data also show that longevity and health are not exclusively affected by DR through the same mechanisms, as there was no statistical correlation between median lifespan and either of our climbing measures. Furthermore, through GWAS across all tested lines, we have implicated a number of new longevity and healthspan genes. In validating three of these genes, we also failed to see a correlation between lifespan and age-related climbing ability. Together these results support the argument that lifespan and healthspan are likely to be regulated by distinct genetic mechanisms.

The separation of lifespan and health regulation has been a topic of debate. Recent studies have suggested either that lifespan and health are uncoupled in their regulation (41, 42, 44, 99) or that they are correlated (71, 100, 101). One recent report demonstrated that known short-lived mutants frequently show a reduction in health by some measures but an extension by others, indicating the complexity of the relationship between lifespan and healthspan (43). It is conceivable that lifespan is uncoupled from healthspan as lifespan is likely to be affected by the weakest link that leads to mortality and thus may not reflect the underlying rate of aging or a particular healthspan trait. Our work here suggests that genotype is a significant contributing factor towards this relationship. One difference between our study and previous reports is that previous work has largely focused on candidate-based targets, whereas we provide the first instance of a direct comparison across a panel of ∼150 strains with naturally arising genetic variation. Through our phenotypic analysis of the DGRP, we saw no correlation between climbing ability and length of life. While climbing ability is certainly not the end-all measure for health, it remains one of the most frequently used and trusted methods for assessing overall functional health (49–52, 54, 102). While others have suggested that functional health by means of physical activity should be presented in the context of maximal functionality rather than absolute activity (101), we further observed no correlation with lifespan in either of these contexts (Fig. 3, Suppl. Fig. 2). Looking further into our DGRP phenotypic data, we did not observe a correlation between lifespan and our functional health measures in general but also saw that the effects of DR were not exclusively beneficial and in some strains affected lifespan and healthspan differentially. Despite the prevailing notion in the field that lifespan-extending interventions will also extend healthspan, our data argues that lifespan and healthspan can be uncoupled quite often. Further, although DR is widely viewed as one of the most robust means for lifespan and healthspan extension (35), our data suggest that genotype significantly influences the extent and type of benefits one can derive from DR. In human studies of longevity and mortality it has been suggested that a healthy lifestyle can improve healthspan without necessarily altering lifespan, creating a compression of the period of disability (103). Understanding the genetic mechanisms which can contribute to compression or extension of morbidity, particularly in response to dietary influences, will allow for more targeted approaches to diagnosing mortality and maximizing healthspan (104, 105). It is worth noting that sex-specific responses to dietary restriction have been observed across different genotypes (31). Furthermore, using walking speed as a predictor of mortality is suggested to be more effective in men than women (45, 106, 107). In our study we use female flies, and we predict that our results could differ from a study conducted entirely with males.

Through GWAS, we were successfully able to pinpoint loci significantly associated with lifespan or climbing regulation. Upon generating a list of statistically significant diet-dependent longevity or health-related loci, we immediately noticed the absence of loci found in genes that take part in the well-studied diet-responsive pathways involved in longevity or health, such as those in the TOR pathway (24). One reason for this could be because polymorphisms that would alter the function of these genes would likely be lethal or inhibit development to adulthood, and thus may not be well-represented in adult wild isolates like the DGRP. A second possibility is that there are many genes which influence lifespan phenotypes and genes in the ILS and TOR pathways represent only a small fraction of those. In support of this argument, over 500 genes have been identified to influence longevity in a variety of models (108), most notably those identified through screens in *S*. *cerevisiae* (109) and *C. elegans* (110, 111).

Through our climbing GWAS, we identified a novel role for the gene *dls* in absolute climbing ability and overall spontaneous activity. Our DGRP strain phenotypes showed several polymorphisms in this gene were associated with the climbing ability only on DR, a diet-specific effect that was mirrored by flies with transposon-based disruption of the *dls* gene. No biological function has been suggested for this gene, and the protein it encodes contains a conserved domain of unknown function (InterPro DUF1091). We have found that expression of this gene increases dramatically on DR in the head, potentially suggesting a neuronal mechanism influencing climbing ability. Interestingly, the longevity effect of a mutation in this gene varied depending on the strain background. We found significantly increased lifespan on both diets in a *w*^*1118*^ background but significantly decreased lifespan on the AL diet in a Canton-S background. Despite these differing results, the significant increases in climbing ability and spontaneous activity on DR were observed in both backgrounds, emphasizing that this gene is not a robust modulator of longevity but appears to regulate overall physical function across multiple genotypes.

One novel longevity locus we identified through our analyses was in *jgh*. The homology of *jgh* to *RGS7BP* in humans suggests a role in regulating neuronal G protein signaling (112). Through our experiments, we verified a diet-specific role in lifespan regulation through *jgh.* Although we observed that knockdown of *jgh* extended lifespan on AL diet, our GWAS association effect was largest on DR. We attribute this difference to the difference between the effects of an intronic single-nucleotide polymorphism in the DGRP strains and RNAi knockdown (113). Human GWAS has previously identified *RGS7BP* as being associated with schizophrenic and bipolar disorders (114) as well as weight gain in response to antipsychotic medication (115), but a dietary link has not been previously observed. Neuronal G protein-coupled receptors have been implicated in *Drosophila* insulin-like signaling (116), providing a potential intriguing, diet-dependent mechanism for further investigation. We also observed a new role for *Fdxh* in diet-responsive longevity, which has previously been shown to regulate mitochondrial function and Friedrich’s ataxia pathology across multiple species (117, 118) and ecdysteroid production in flies (87). Studies in multiple model systems (119–121) and humans (122) have shown the importance of proper Fe-S maintenance and the role of these clusters in proper electron transport in the mitochondria. Given the role of mitochondrial function in diet-dependent effects on longevity (23, 123–126), it is fitting that our screen revealed a role for a mitochondrial gene in lifespan regulation. Additionally, *Fdxh* has been observed in an array for cycling circadian genes in the head (127). As circadian clocks have been implicated in diet-dependent lifespan extension (4), modulation of circadian phenotypes is another potential mechanism by which *Fdxh* could modulate diet-specific longevity. One human homolog of *Fdxh, FDX1L*, has been implicated in inflammatory bowel disease and Crohn’s disease (128) and dermatitis (129). Another more distantly related human homolog, *COX15*, has been found in GWAS for childhood obesity (130), Crohn’s disease (131), colorectal cancer (132), and cardiovascular disease (133), all of which could provide clear impacts on longevity.

Together, we have provided a novel approach to understanding the natural genetic factors which regulate diet-dependent changes in longevity and health. By measuring both lifespan and also an age-related component of health, climbing ability in the same strains, we were able to dissect the genetics of two age-related traits simultaneously. Our experiments, using diet manipulation, have further detailed the diversity of responses in wild strains to DR, which varies greatly by genotype both in lifespan and climbing ability. Most previous studies have examined healthspan in known longevity genes, which may be subject to confirmation bias and negative results where lack of correlation between healthspan and lifespan was obtained but under-reported. Our study utilized an unbiased approach to examine the relationship between healthspan and lifespan in over 150 strains and manipulation of candidate genes identified from this analysis. Our findings strongly argue for genetic uncoupling of mechanisms that modulate longevity and healthspan. The independence of healthspan from lifespan may have important bearing in designing dietary interventions that delay the effects of aging in humans and other species.

## Materials and Methods

### Fly lifespan phenotyping

DGRP lines were obtained from Bloomington Stock Center, Bloomington, IN (134). Each line was mated and developed on a standard lab diet (1.5% yeast). Two to three days post-eclosion, mated female progeny were transferred to AL (5.0% yeast extract) or DR (0.5% yeast extract) diet, as previously described (23, 135). Eight vials of 25 flies were used per diet per strain. Flies were maintained at 25°C and 65% relative humidity throughout life. Living flies were transferred to fresh vials every other day, with dead flies being recorded, until all flies were dead. One biological replicate (200 animals) was recorded for 107 lines, two biological replicates for 52 other lines, and three biological replicates for two other lines. *w*^*1118*^ was also tested with each batch as an internal control. DGRP lines not tested were not viable long term in our lab.

### Fly climbing phenotyping

Throughout life, climbing ability was recorded weekly on days between vial transfers for all vials containing 20 or more living flies. The negative geotaxis climbing ability test was adapted from previous methods (136). Flies were placed in an empty vial with a line 6 cm from the bottom. Flies were gently tapped to the bottom of the vial and the number able to cross the line within 10 seconds was recorded. This was repeated three times for each vial, and the percentage of live flies still climbing above the line was averaged for a given line at weekly timepoints throughout life. For normalized climbing values, weekly climbing values were normalized to the percentage of flies climbing one week following placement on AL or DR. We used the day at which flies passed below 50% of their day seven climbing value for genome-wide analysis, as well as the day at which a 20% or less of a surviving population of flies were still able to climb.

### Genome-wide association analysis

We used DGRP release 2 genotypes, and FlyBase R5 coordinates for gene models. As in Nelson *et al.*, 2016 (77), we used only homozygous positions and a minor allele frequency of ≥25% to ensure that the minor allele was represented by many observations at a given polymorphic locus. The collected phenotype and genotype data were used as input into an association test via ordinary least squares regression using the StatsModels module in Python (137). The linear model was phenotype = β_1_ x genotype + β_2_ x diet + β_3_ x genotype x diet + intercept. Nominal *p*-values denoted as “genotype” in Table 1 report the probability that β_1_?≠?0, and those denoted as “interaction” report the probability that β_3_?≠?0. For the binary “case-control” search for determinants of long lifespan, we used median lifespan thresholds of ≥41 days on AL diet and ≥51 days on DR diet, and a Fisher’s exact test comparing the long-lived and short-lived populations with both alleles at a given position. To avoid the potential for false positives at a given nominal cutoff owing to *p*-value inflation, we calculated false discovery rates via permutation as follows: for a given permutation *i*, we randomized phenotype values across DGRP lines, retaining the true diet assignment, and on this permuted data set we carried out association tests for each marker in turn as above. We counted the number of markers *n^i^* that scored above a given *p*-value threshold *t*. We tabulated the false discovery rate (FDR) at *t* as the ratio between the average *n^i^* across ten permutations and the number of markers called at *t* in the real data. We used an empirical FDR upper bound of 10% within a given analysis to call candidate loci of interest.

### Gene expression analysis

To determine gene expression in a normal system, we sampled five whole flies, 50 heads, 50 thoraxes, or 50 abdomens from mated females *of w^1118^* control strain after one week on AL or DR diet. We isolated RNA using Zymo Quick RNA MiniPrep kit (R1054) (Zymo Research, Irvine, CA). For qRT-PCR, we used Superscript III Platinum SYBR Green One-Step qRT-PCR kit from Invitrogen, Carslbad, CA (11736-051) and followed manufacturer’s instructions with a Roche Lightcycler 480 II machine. To validate the effects of RNAi or mutation on gene expression, we collected five whole female bodies or 50 heads following one week on AL or DR. We then isolated RNA from these samples and performed qRT-PCR on the perturbed genes as described.

### Gene alteration phenotyping

For candidate gene validation, all lines were obtained from Bloomington Stock Center (134) or Vienna *Drosophila* RNAi Center, Vienna, Austria (138) (see Suppl. Fig. 9 for list of lines used (87, 139–148)). To validate the GWAS-predicted effects of *jgh* and *Fdxh*, we used the whole-body GeneSwitch (149) driver *Act5C*-GS-Gal4 (140) and the neuron-specific driver *Elav*-GS-Gal4 (139) for directed RNAi. 15 virgin driver females were mated with three transgene line males in four bottles containing a standard diet. Two-three days following progeny eclosion, mated females were sorted onto AL or DR media with or without 200 mM RU486 (final concentration) for RNAi activation (4, 135), and flies were maintained on these media for life. For *dls* analysis, a *Minos* element mutant line was used for gene disruption (85). This line was outcrossed to *w*^*1118*^ or Canton-S control strains for six generations using a GFP tag associated with the inserted element (144). Spontaneous activity was measured on day 5 after flies were placed on either the AL or DR diet. Three vials of 25 female flies for each condition were placed for 48 hours in a 12-hour light-dark cycle at 25°C and 65% relative humidity in a TriKinetics Drosophila Activity Monitor system (TriKinetics, Waltham, MA), and beam crosses were recorded for 24 hours (150). Activity was recorded for three separate biological replicates.

## ACKNOWLEDGEMENTS

K.A.W. is supported by NIH/NIA F31 award AG052299. C.S.N. was supported by NIH/NIA F32 award AG047024. This work was funded by grants from the American Federation of Aging Research (R.B.B. and P.K.), NIH grants R01AG038688 and AG045835 (to P.K.) and R01AG049494 (to Dr. Daniel Promislow) and the Larry L. Hillblom Foundation. We thank the Bloomington Drosophila Stock Center and the Vienna Drosophila Stock Center for flies. We thank the members of the Kapahi lab for helpful discussions, as well as Dr. Daniel Promislow, Dr. John Newman, and Kelly Jin for their feedback.

## AUTHOR CONTRIBUTIONS

K.A.W., C.S.N., R.B.B., and P.K. designed research; K.A.W.,C.S.N., and J.N.B. performed research; K.A.W., C.S.N., R.B.B., and P.K. analyzed data; K.A.W., C.S.N., and P.K. wrote the paper.

